# CaV1.3 enhanced store operated calcium promotes resistance to androgen deprivation in prostate cancer

**DOI:** 10.1101/2021.09.03.458558

**Authors:** Debbie O’Reilly, Tim Downing, Sana Kouba, Marie Potier-Cartereau, Declan J McKenna, Christophe Vandier, Paul Buchanan

## Abstract

Androgen deprivation therapy (ADT) is the main treatment for advanced prostate cancer (PCa) but resistance results in progression to terminal castrate resistant PCa (CRPC), where there is an unmet therapeutic need. Aberrant intracellular calcium (Ca_i_^2+^) is known to promote neoplastic transformation and treatment resistance. There is growing evidence that expression of voltage gated calcium channels (VGCC) is increased in cancer, particularly the *CACNA1D*/CaV1.3 in CRPC. The aim of this study was to investigate if increased CaV1.3 drives resistance to ADT and determine its associated impact on Ca_i_^2+^ and cancer biology.

Bioinformatic analysis revealed that *CACNA1D* gene expression is increased in ADT treated PCa patients regardless of TMPRSS2:ERG status. Corroborated in both in vivo LNCaP xenograft mouse and in vitro PCa cell line models which demonstrated a significant increase in CaV1.3 protein expression following ADT with bicalutamide. The expression was found to be a shortened 170kDA CaV1.3 isoform associated which failed to mediate calcium influx following membrane depolarisation. Instead, under ADT CaV1.3 mediated a rise in basal cytosolic calcium and an increase in store operated calcium entry (SOCE). This in turn drove both proliferation and survival of long-term ADT CRPC cells.

Overall, this study demonstrates for the first time in PCa that increased SOCE through a novel CaV1.3 mechanism which represents a novel oncogenic switch that contributes to ADT resistance and promotes CRPC biology. Highlighting aberrant intracellular calcium in CRPC as a potential area for therapeutic development to improve patient outcomes.

## Introduction

Worldwide prostate cancer (PCa) is the most frequently diagnosed malignant tumour in men over 60 years old^1^. Androgen deprivation therapy (ADT) is the main treatment for advance or metastatic disease but associated relapse results in the development of terminal castrate resistant prostate cancer (CRPC)^2^. Further understanding into the mechanisms driving androgen-independent growth is required to highlight new therapeutic targets and improve patient outcomes.

Intracellular calcium (Ca_i_^2+^) modulation regulates many key cellular processes such as migration, proliferation, apoptosis and expression^3^. Its disruption is known to promote the development of various cancer hallmarks and drive disease progression^4^. The regulation of Ca_i_^2+^ is controlled by a range of calcium channel families, the expression of which is altered across various malignancies^4^, including PCa^5,6^. In non-excitable cells, such as that found in cancer, Ca_i_^2+^ is modulated through store operated current (SOC)^7^, mediated by release from intracellular stores and associated store operated calcium entry (SOCE)^7^. Research shows that aberrant SOC is linked to various aspects of cancer progression and treatment resistance^8^. Ca_i_^2+^ can also be modulated by membrane potential regulating voltage gated calcium channels (VGCC)^9^, but this is normally found in excitable cells. However, microarray meta-analysis of cancer samples has highlighted increases in VGCC gene expression across a range of cancer types^10^. Furthermore, emerging studies show that this family can contribute to aberrant Ca_i_^2+^ and drive cancer biology^4,11,12^.

There is growing evidence that gene expression of L-type VGCC,*CACNA1D*, is upregulated in PCa tissues compared to normal^6^ and that it is associated with Gleason score and lymph node status^13^. In addition, it is a known target of the common somatic alternation present in 50% of PCa patients, TMPRSS2:ERG^14^. To date, the single study in PCa exploring its protein form, CaV1.3, found it increased Ca_i_^2+^ following androgen stimulation of androgen receptor (AR) positive PCa cell lines^15^. Channel inhibition with calcium channel blockers (CCB), lead to reduced cell proliferation^15^. Similar observations were also made in studies on breast and endometrial cancer cells^16,17^. However, CaV1.3’s mechanism of action has not been explored in PCa. Traditionally it promotes calcium influx following depolarisation but there is growing evidence of potential non-canonical roles through different channel isoforms^18^. Research previously published by this group in colorectal cancer, demonstrated that CaV1.3 could promote an increase in SOC through interaction with the sodium calcium exchanger^19^. In addition, its outlined role in mediating androgen stimulated Ca_i_^2+^ also suggests an association with SOC, due to its known activation downstream of G-protein coupled receptors such as AR^20^. Interestingly, in PCa ADT is known to contribute to aberrant SOC which in turn supports the development of CRPC^21^. Linked to this, *CACNA1D* gene and protein (CaV1.3) expression has been found to be further increased in CRPC patient samples^13,15^. Despite this the impact of CaV1.3 on Ca_i_^2+^ and associated biology under ADT has not been explored.

Taken together, this suggests that the upregulation of CaV1.3 on progression to CRPC drives aberrant SOC resulting in the development of ADT resistance. This study aimed to explore for the first time the link between ADT and CaV1.3 expression, as well as how this treatment affected the channels functional ability to influence Ca_i_^2+^ and PCa biology. Our novel study shows in both in vivo and in vitro models that that enhanced CaV1.3 expression is a significant contributing factor to developing ADT resistance and progression to CRPC. Furthermore, this change was shown to enhanced Ca_i_^2+^ through a SOCE mechanism which in turn promoted proliferation and survival in CRPC.

## Materials and Methods

### Cell Culture

Human Prostate cancer cell lines used include, AR positive androgen sensitive LNCaP (ATCC # CRL-1740), androgen resistant CRPC LNCaP-Abl^22^ and intermediate ten-day ADT, LNCaP-ADT. All were maintained in RPMI (Gibco) with 10% fetal bovine serum (Gibco) in a humidified incubator at 37°C and 5% CO2. LNCaP-Abl and LNCAP-ADT where kept under ADT with 10µM bicalutamide (Sigma).

### Prostate xenograft Mouse model

Mice tumour tissue was obtained from previously generated LNCaP xenografted BALB/c immune-compromised (SCID) mice^23^. Tumours were established by subcutaneous injection of LNCaP cells (2×10^6^ suspended in 100 μl of Matrigel) followed by daily treatment administered orally (p.o.) over the course of 21 days with either bicalutamide prepared in vehicle (0.1% DMSO in corn oil) or vehicle only. Frozen tissue samples were slowly thawed over ice and sectioned with a sterile blade.

### PCR

Isolation of cell line RNA was conducted using the High Pure RNA isolation kit (Roche). RNA was isolated from mouse xenograft tumour tissue following homogenisation with a sterile pipette tip in TRIzol reagent (Invitrogen). Total RNA quantified with a nanodrop and purity determined through A_260_/A_280_ ratio. cDNA was synthesised according to instructions for the Transcriptor first strand cDNA synthesis kit (Roche). Lyophilised primers (Sigma Aldrich) were designed using exon spanning sequences in primer3. Gene levels determined using Faststart essential Syber Green master mix (Roche) on a Roche Lightcycler Nano, reaction mix as instructed with 0.5uM forward and reverse primer and 50ng cDNA. The housekeeping gene HPRT1 was used to calculate relative gene expression with the -ΔΔCT method.

### Protein preparation

Total cell lysate was extracted from ice cold cells using RIPA (Thermo Scientific) plus Halt inhibitor cocktail (Thermo Scientific). A cell fractionation kit (Abcam) was used, extracting three fractions, the cytosolic, nuclear and membrane fraction (membrane organelles and the plasma membrane). The included manufactures protocol was followed using approximately 6×10^6^ cells for each sample, which were quantified using a Pierce BCA Protein Assay Kit (Thermo Scientific).

### Western Blot

Electrophoresis was performed using the Mini Gel Tank (Invitrogen) and 4-12% Bolt Bis-Tris Plus pre-cast gradient gels (Invitrogen). 50ug of protein per lane was run and transferred using the Mini Blot module at 20V for 60 minutes onto 0.4µm polyvinylidene fluoride (PVDF) in recommended buffers (Invitrogen). Membranes were blocked for one hour at room temperature (RT) in 5% milk powder TBS-T (Tris buffered saline with tween 20 (0.1%)). Primary antibody (CaV1.3, Abcam, mouse monoclonal, AB-84811) was used 1:500 in 5% milk TBS/T overnight at 4°C. Secondary antibody incubated for one hour at RT in 5% milk TBS-T at 1:1,000 (HRP-Anti-Mouse, BD Pharmigen, Goat Polycloncal, 554002). Between steps 3 × 5 minute washes were performed in TBST. The membrane was incubated with Supersignal West Dura Chemiluminescent Substrate (Thermo Scientific) and exposed to X-ray film before development. Band densitometry conducted using Image J relative to housekeeping protein Actin (1:10,000, BD Transduction laboratories, Mouse Monoclonal, 612656).

### siRNA

Cells were seeded in plates and grown to 80% confluency in Opti-MEM (Gibco). Dharmacon transfection reagent was used alongside Dharmacon ON-TARGETplus siRNA for Non-targeting (Negative control, sictr) and *CACNA1D* siRNA (SiCaV1.3). The transfection siRNA mixture was generated with the appropriate volumes in the manufacturer’s handbook. Cells where treated with transfection mix in Opti-MEM for 48-hours prior to RNA extraction or Ca^2+^ imaging, or 72-hours prior to protein extraction. Percentage knockdown was determined (Figure S3), achieving >70% in gene and >80% in protein knockdown across all three cell types.

### Cell Proliferation

Cell proliferation was measured by adding WST-1(Roche, UK) as detailed in manufactures’ literature and incubating for 4 hours. Afterwards colorimetric readings at 450nm and 690nm (Background) were made using the VICTOR multilabel plate reader. The background control was subtracted from each measurement and triplicate average used in the analysis.

### Colony Formation Assay (CFA)

PCa cells seeded at a density of 2500 cells per well and treated 48 hours later. After 14 days of growth the media was aspirated off, cells were washed in PBS before fixing in methanol for 30 minutes followed by staining in 1:10 crystal violet solution for 1 hour. After washing in distilled water, a collection of 50 or more cells was considered a colony and counted.

### Migration

The Xcelligence™ system, Boyden chamber-based assay was used to determine cellular migration. Pre-treated cells were seeded at a density of 8×10^4^ cells/well in duplicate in the top chamber of the cell invasion and migration (CIM) plate in 100µl media containing 2% FBS. The bottom chamber had media containing 10% FBS as a chemoattractant or 2% FBS in the negative control wells. CIM plates were set into the Xcelligence™ contained within the incubator at 37°C with 5% CO2. The cell electrical impedance was recorded every 15mins for 60 hours.

### Calcium measurements

To determine relative changes in cytosolic calcium (Ca_c_^2+^), PCa cells were suspended in OptiMEM (Gibco) and loaded with 2μM Fura-2-AM ratiometric dye diluted in DMSO in the dark for 45min. Cells were washed twice in OptiMEM and left for 30min, before being resuspended in Ca^2+^free physiological saline solution (PSS) and seeded into a 96 well black plate. Florescence was measured at 510nM after excitation at 340 and 380nM using a VICTOR multilabel plate reader. SOC measurements were recorded over time after 5µM Thapsigargin induced endoplasmic reticulum depletion followed by application of 2mM CaCl_2_ to measure Ca^2+^ entry (SOCE). Membrane depolarisation was induced by addition of high external potassium solutions of 56 mM KCl (K60) or 76 mM KCl (K80), alongside osmolality controls of 56 mM NaCl (Na196) or 76 mM NaCl (Na216). To ensure CaV1.3 was not already inactivated cells were first hyperpolarised with potassium channel blocker NS1189. PSS (in mM): NaCl 140, MgCl_2_ 1, KCl 4, CaCl_2_ 2, D-glucose 11.1 and HEPES 10, adjusted to pH 7.4 with NaOH. The Ca^2+^ free solution or 0Ca is a PSS solution without CaCl_2_ and with 1 mM EGTA. Calcium measurement of peak amplitude of Tg response and Ca^2+^ entry was determined by calculating the ratio between baseline and the maximal Ca^2+^ after application of Tg or 2mM Ca^2+^. The slope for both was determined by linear regression curve fitting 20 sec after treatment to estimate the speed of movement. Area under the curve for total SOC was calculated using Graphpad prism 5 with the trapezoid rule.

### Prostate Cancer Patient Bioinformatics

#### Data retrieval, quality control and normalisation

DNA Affymetrix Human Exon 1.0 ST microarray data^24^(n=185 samples, GSE21032) with 6,553,600 potential probes was downloaded with R packages Biobase v2.30.0 and GEOquery v2.40.0. After quality control (QC), this had information for n=130 primary cancers, n=18 metastases and n=24 control normal adjacent epithelial samples (Table S1). 29 of the primary and adjacent samples were paired. Cell line or xenograft samples were excluded. QC and visualisation with principal components analysis (PCA) was implemented with Bioconductor v3.9 to check for outliers based on PCA, which excluded PAN136, PCA040 and PCA208. The data was background-adjusted, quantile-normalised, and log-transformed using the RMA algorithm in affy v3.0^25^ to obtain the samples’ normalised expression across 22,011 informative probes.

#### Differential expression testing and comparison with patient phenotypes

Differential gene expression testing was implementing using limma v3.34.4 based on linear models of the log_2_ fold-change (LogFC) and adjusted p value. Statistical analyses and comparisons with phenotypes were completed with R packages dplyr v0.7.6, ggplot2 v3.0.0, grid v3.4.1, magrittr v1.5, plyr v1.8.4, readr v1.1.1, survival v2.42.6, tidyr v0.8.1, tidyverse v1.2.1, tidyselect v0.2.4, and xtable v1.8.2. 124 raw p values were corrected for multiple tests using the Benjamini-Hochberg approach.

### Statistical analysis

All graphs were prepared using Prism (Graphpad software, USA). Results are expressed as mean+/-s.e.m unless otherwise stated. Parametric tests were employed when N>10 and included two-tailed unpaired t-test for comparison between two samples, one-way ANOVA with Tukey multiple comparison test (MCT) between 3+ groups and two-way ANOVA with Tukey MCT for differences between cell types that have been treated with or without siRNA. Non-parametric tests were used for non-normal sample distributions and N<10. Non-parametric tests between two groups used a Mann-Whitney test or for multiple groups a Kruskal-Wallis with Dunn’s MCT. The statistical tests employed are described in the figure legend, with annotations as *P<0.05, **p<0.01, ***p<0.001. All results are generated from at least three independent experiments denoted by the N number, with the total number of individual repeats within each denoted by the n number.

## Results

### Increased *CACNA1D* expression with PCa progression is associated with ADT

We tested if *CACNA1D* gene expression was positively correlated with PCa stage and other clinical parameters (Table S1). Differential expression testing of existing microarray data^24^, showed that *CACNA1D* expression was 11.3% higher on average in metastatic compared to primary tumours (p=0.035, Figure 1A). The 28 paired primary samples had 10.3% higher *CACNA1D* than their adjacent normal pairs (p=0.018). Similarly, the metastatic tumours had 23.5% higher *CACNA1D* expression than the adjacent normal samples (p=1.14×10^−7^). *CACNA1D* expression was higher in metastatic CRPC, so we tested if the development of resistance to ADT was a driver of the increase in this terminal group. We found that ongoing ADT samples had higher expression than the untreated (p=0.041), as did the pooled ongoing ADT and post-ADT relative to the untreated group (p=0.045, Figure 1B). Higher *CACNA1D* expression was associated with an elevated combined Gleason score in the combined primary and metastatic tumour samples (Figure 1C). This was supported by observations of higher *CACNA1D* expression in primary samples with Gleason score 7 relative to 6 (n=41 vs n=73, t-test p=7.0×10^−3^), across Gleason scores 6 to 8 for all samples (one-way ANOVA p=0.012) and across the primary ones alone (one-way ANOVA p=0.018).

**Figure 1.**
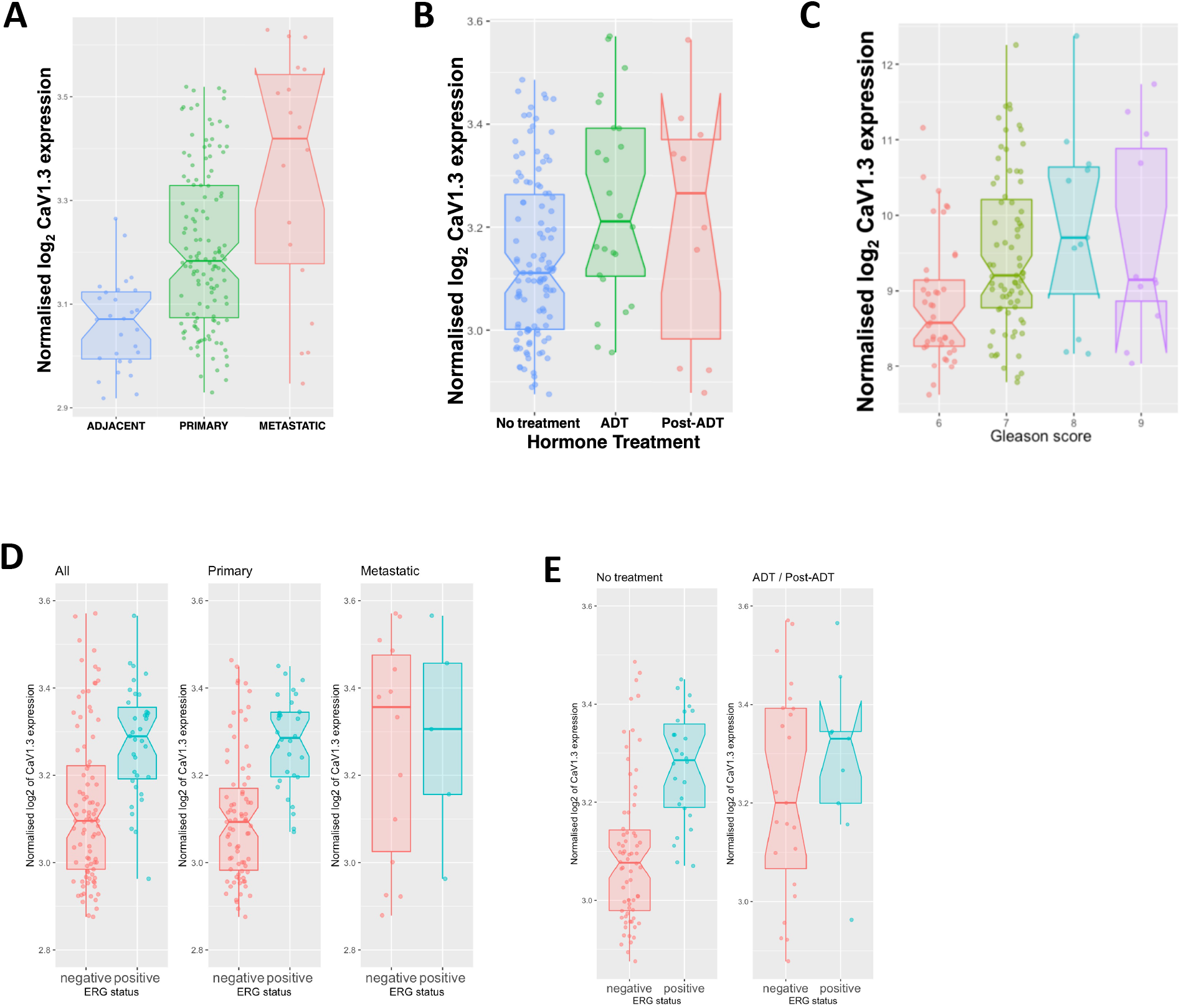
Prostate cancer patients demonstrate an increase in *CACNA1D* (CaV1.3) expression associated with sample type, androgen deprivation treatment, *ERG* status and Gleason score. **(A)** Normalised *CACNA1D* (y-axis) expression was higher in the metastatic (red, N=18) than the primary (green, n=130) tumours, and both were elevated compared to the adjacent normal samples (blue, N=28) (t-test). **(B)** *CACNA1D* expression was higher in the post-ADT (red, N=10, median=3.27) than the ongoing ADT (green, N=24, median=3.21) samples, and both were elevated invidually or pooled (N=34) compared to the untreated (blue, N=114, median=3.11)(t-test). **(C)** *CACNA1D* expression was positively correlated with a higher Combined Gleason score for the combined primary and metastatic samples (t-test and one-way ANOVA). **(D)** *CACNA1D* expression was higher in *ERG*-positive (red) than *ERG*-negative (blue) tumours in the combined (left panel, n=148), the primary (middle panel, n=130) but not the metastatic (right panel, N=18) samples (t-test). **(E)** *CACNA1D* expression (left panel) was higher in the untreated *ERG*-positive (blue, N=26) than the untreated *ERG*-negative (red, N=70) patients. *CACNA1D* expression in hormone-treated patients (right panel) was marginally but not statistically higher in *ERG*-positive (blue, N=9) than *ERG*-negative (red, N=23) patients (t-test). Across all the median values are shown by the horizontal bars, and the notch reflect the interquartile range scaled by the sample size.

The somatic alteration, TMPRSS2:ERG, is present in about half of all PCa cases resulting in ERG upregulation^14^. *CACNA1D* is a known ERG target^15,26–29^ and here its expression was associated with ERG status. Across all samples, those with an ERG-positive status (n=34) had higher *CACNA1D* levels than ERG-negative (n=91) or flat (n=22) profiles (t-test p=1.46×10^−5^, Figure 1D). This association held in the primary samples, were ERG-positive samples (n=30) had higher expression than ERG-negative (n=78) or flat (n=22) profiles (t-test p=1.04×10^−6^), but not for metastatic samples (n=14 ERG-negative vs n=4 ERG-positive), where expression was elevated in both groups (Figure 1D). Metastatic disease is typically associated with ongoing or post ADT, we sought to explore further if ADT treatment had an effect on *CACNA1D* expression independent of ERG regulation (Figure 1E). Here we observed untreated ERG-positive patients had higher CaV1.3 expression than untreated ERG-negative patients (p=2×10^−7^) (Figure 1E). In contrast, patients who did receive ADT had elevated *CACNA1D* expression in both ERG-Negative and ERG-Positive groups (Figure 1E). Thus in ERG-negative samples, CaV1.3 expression was elevated in patients receiving hormone therapy compared to those who did not (p=0.0106).

### ADT enhances CaV1.3 expression in PCa cell line and mouse models

Using a cell line model of PCa disease progression, we aimed to validate this observed increase in CANCA1D expression with disease progression and ADT (Figure 1). We found that *CACNA1D* gene expression was increased in short term ADT treated, LNCaP-ADT (1.7FC+/-0.13sem, p<0.0003, Figure 2A). While CaV1.3 protein expression in the same model, showed a significant increase of 2.77FC+/-0.2sem in long term ADT CRPC, LNCaP-Abl(p<0.027, Figure 2B) but no change in LNCaP-ADT.

**Figure 2:**
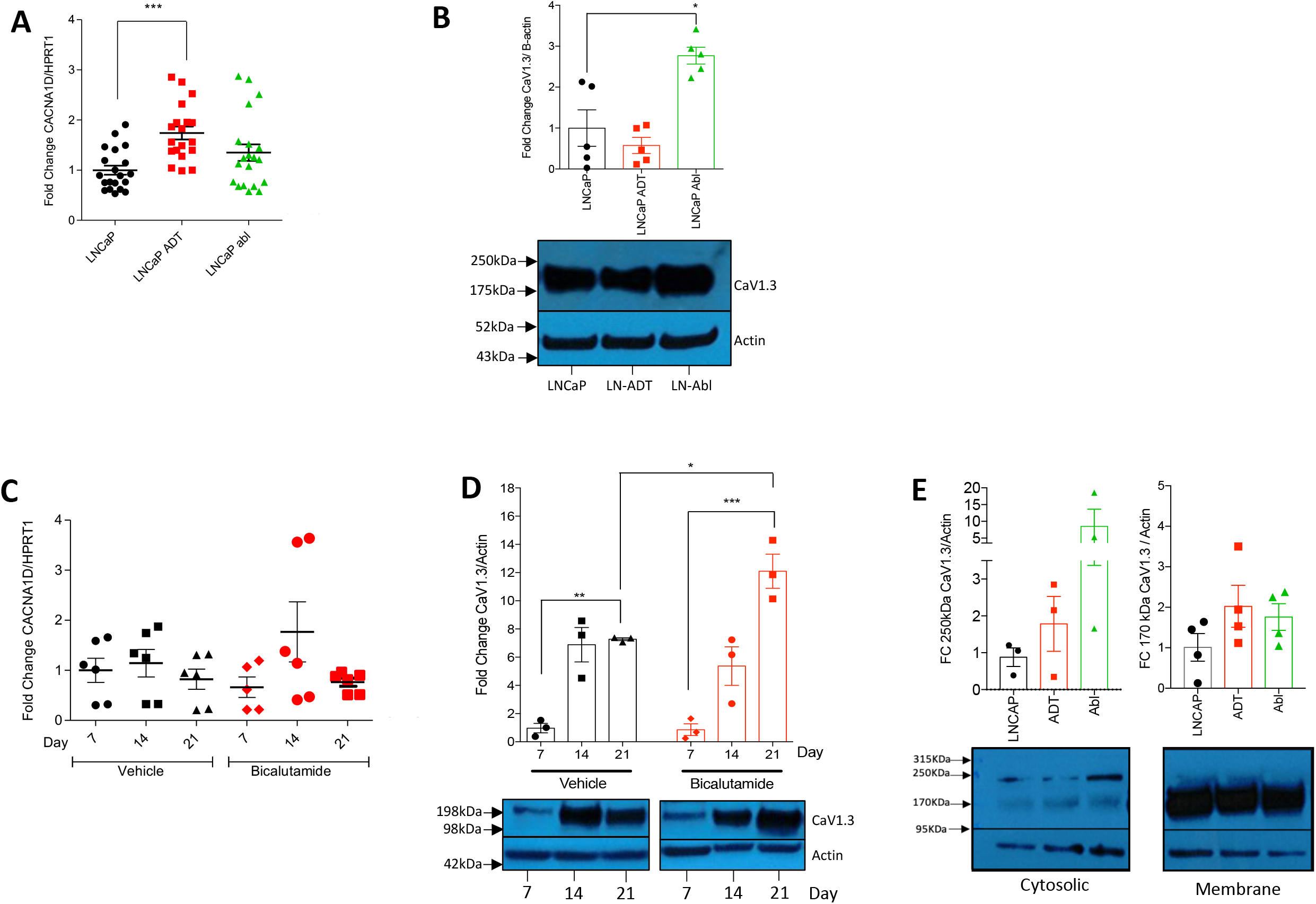
Expression of CaV1.3 is increased after ADT with bicalutamide in both in vitro PCa cell line and in vivo xenograft mice models. **(A)** *CACNA1D* gene expression was measured by qPCR and fold change calculated compared to housekeeping HPRT1 in untreated LNCaP, short term ADT (LNCaP-ADT) or long term ADT (LNCaP-Abl) and normalised to LNCaP (N=10, n=20, Kruskal-Wallis-Dunn’s MCT). **(B)** CaV1.3 protein expression in the outlined PCa cell lines was determined by western blot. Fold change was measured by densitometry, normalised to LNCaP and displayed in a bar graph, with a representative blot shown in the lower panel. (N=5, Kruskal-Wallis-Dunn’s MCT). **(C)** Gene expression of *CACNA1D* was determined in xenografted LNCaP tumours grown in BALB/c mice by qPCR, treated daily with either 10µM/2kg bicalutamide (Red) or equivalent vehicle control (Black) with samples taken every 7 days up to 21 days. Fold change calculated normalised to corresponding day 7 control and displayed on bar graph. (N=3, n=6, Kruskal-Wallis-Dunn’s MCT). **(D)** CaV1.3 expression in outlined PCa mouse model was determined by western blot. Fold change was measured following quantification by densitometry normalised to day 7 vehicle and displayed as a bar chart, with a representative blot displayed below (N=3, two-way ANOVA, Tukey’s and Bonferroni’s MCT either within or across treatments respectively). **(E)** PCa cell line protein was separated into cytosolic (N=3) and membranous fractions (N=4) with CaV1.3 fold change expression calculated by densitometry following western blot with each normalised to LNCaP (Kruskal-Wallis-Dunn’s MCT). All PCa cell lines represented as the following, Black circle - LNCaP, Red square – LNCaP-ADT, Green Triangle – LNCaP-Abl.

To further confirm the link between *CACNA1D*/CaV1.3 expression and ADT resistance, a bicalutamide-treated LNCaP Xenograft mouse model was employed. There were no differences in *CACNA1D* gene expression across any of the three time points in either bicalutamide-treated or vehicle control samples (Figure 2C). However, CaV1.3 protein expression increased over time with tumour growth in both vehicle and ADT samples (p<0.0001). Culminating in an increase in ADT (12.09FC+/-1.2sem, p<0.0001) and vehicle (7.27FC+/-0.1sem, p=0.0063) treated groups compared to their corresponding day 7 controls (Figure 2D i+ii). Overall, CaV1.3 expression was 66.29% higher in the ADT group at day 21 (p<0.0092) compared to its associated day 21 vehicle control (Figure 2D).

CaV1.3 protein detected in both cell line and mouse models had a molecular weight of ∼170kDa (Figure 2B and D). To determine the cellular location and the potential presence of other isoforms in our PCa cell line model, protein fractionation was performed. Here again the 170kDa variant was the most prominently expressed, appearing in the membranous fraction, which includes the plasma membrane and intracellular organelles. Here its expression was increased in both ADT (2.0FC+/-0.5sem) and Abl (1.76FC+/-0.3sem). In the cytosolic fraction, expression of the 250kDa isoform was detected, with increased expression in LNCaP-ADT (1.78FC+/-0.7Sem) which was further upregulated in LNCaP-Abl (8.5FC+/-5.1sem).

### Increased CaV1.3 under ADT corresponds with increased basal cytosolic calcium

Owing to the observed upregulation of CaV1.3 under ADT we sought to determine its role in modulating Ca_i_^2+^. To test CaV1.3 voltage gating, increasing concentrations of high external potassium (60 and 80mM) were added to induce membrane depolarisation but neither induced any change in Ca_i_^2+^, either with or without ADT (Figure 3A and 3Bi+ii). High sodium controls (60 and 80mM) (Figure 3A and 3Biii), showed no effect on Ca_i_^2+^ measurements, highlighting that the above result was not influenced by any changes in osmolality due to high external potassium. To investigate further basal calcium levels were investigated. Total basal Ca^2+^ was measured in the presence of 2mM external Ca^2+^ (Figure 3C i+ii) and no significant increase was detected across all three cell lines. However, basal Ca_c_^2+^ concentration measured in 0mM external Ca^2+^ (Figure 3Di+ii), registered a F340/F380 increase in LNCaP-Abl of 1.16 (+/-0.03sem, p<0.0019) and LNCaP ADT to 1.065 (+/-0.03sem), both compared to LNCaP (1 +/-0.03sem).

**Figure 3:**
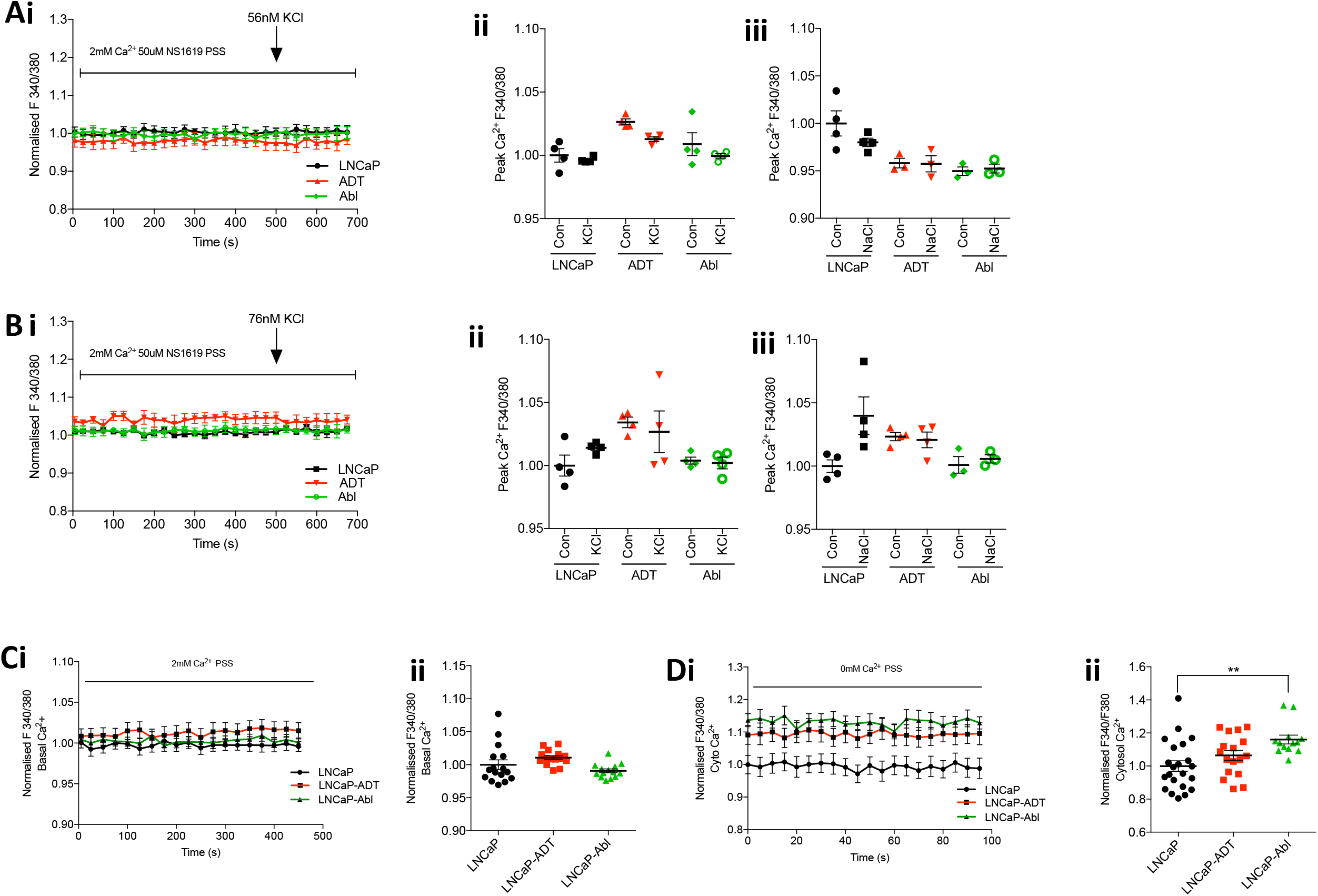
Increased CaV1.3 under ADT influences basal cytosolic calcium through a non-canonical mechanism. PCa cells were loaded with Fura 2-AM for ratiometeric analysis of calcium (Ca2+). Canonical CaV1.3 voltage gated function was tested by depolarisation with addition of high external potassium of **(A)** 56nM KCl (N=4) or **(B)** 76nM KCl (N=4) and represented as **(i)** mean calcium trace over time or **(ii)** peak fold change after treatment normalised to LNCaP (Kruskal-Wallis-Dunn’s MCT). Matched osmolarity controls **(iii)** of 56 mM or 76mM NaCl displayed as peak fold change after treatment normalised to LNCaP (Kruskal-Wallis-Dunn’s MCT). **(C)** Basal calcium was measured (N=4) in 2mM external Ca^2+^ for LNCaP (n=16), LNCaP-ADT (N=15) or LNCaP-Abl (n=14) and **(D)** Cytosolic calcium (N=8) was measured in 0mM external Ca^2+^ for LNCaP (n=22), LNCaP-ADT (N=17) or LNCaP-Abl (n=13), each are displayed as **(i)** trace over time or **(ii)** peak fold change in basal calcium (One-way ANOVA, Dunnett’s).

### Enhanced SOCE under androgen deprivation conditions is modulated through CaV1.3

The observed increase in basal Ca_c_^2+^ and CaV1.3 expression, suggests an associated link to SOC. To investigate further, intracellular store release and associated Ca^2+^ entry were measured either with or without siRNA CaV1.3 knockdown. Validation of which observed a 70% and 80% reduction in both gene and protein levels respectively on average across all three cell lines (Figure S3). In relation to intracellular store release, Tg produced a F340/F380 ratio increase in calcium across all three cell lines under non-targeting siRNA conditions, with LNCaP (1.14+/-0.02sem), LNCAP-ADT(1.15+/0.02sem) and LNCaP-Abl(1.34+/-0.02sem) (Figure 4A, p<0.0001, at 275sec). No difference was observed in Tg response following CaV1.3 siRNA targeting in LNCaP (1.17+/-0.02sem), LNCaP-ADT (1.13+/-0.02sem) and LNCaP-Abl (1.37+/-0.04sem) (Figure 4B). Normalisation of these results to LNCaP sictr showed that LNCaP-Abl produced a Tg induced Ca^2+^peak of 1.9FC+/-0.09sem and 1.9FC+/-0.2sem under sictr and SiCaV respectively (p<0.0001, Figure 4C). Two-way ANOVA analysis confirmed that ADT treatment affected store release (p<0.0001) but CaV1.3 did not. Analysis of Tg-induced rising Ca^2+^ slope demonstrated a similar effect to Tg peak, showing no difference between LNCaP and LNCaP-ADT either under sictr or siCaV conditions but with LNCaP-Abl demonstrated an increase of 2.03+/-0.21sem (p<0.0263) and 2.41+/-0.42sem (p<0.0006) respectively, again due to ADT (p<0.0001) but not CaV1.3.

**Figure 4:**
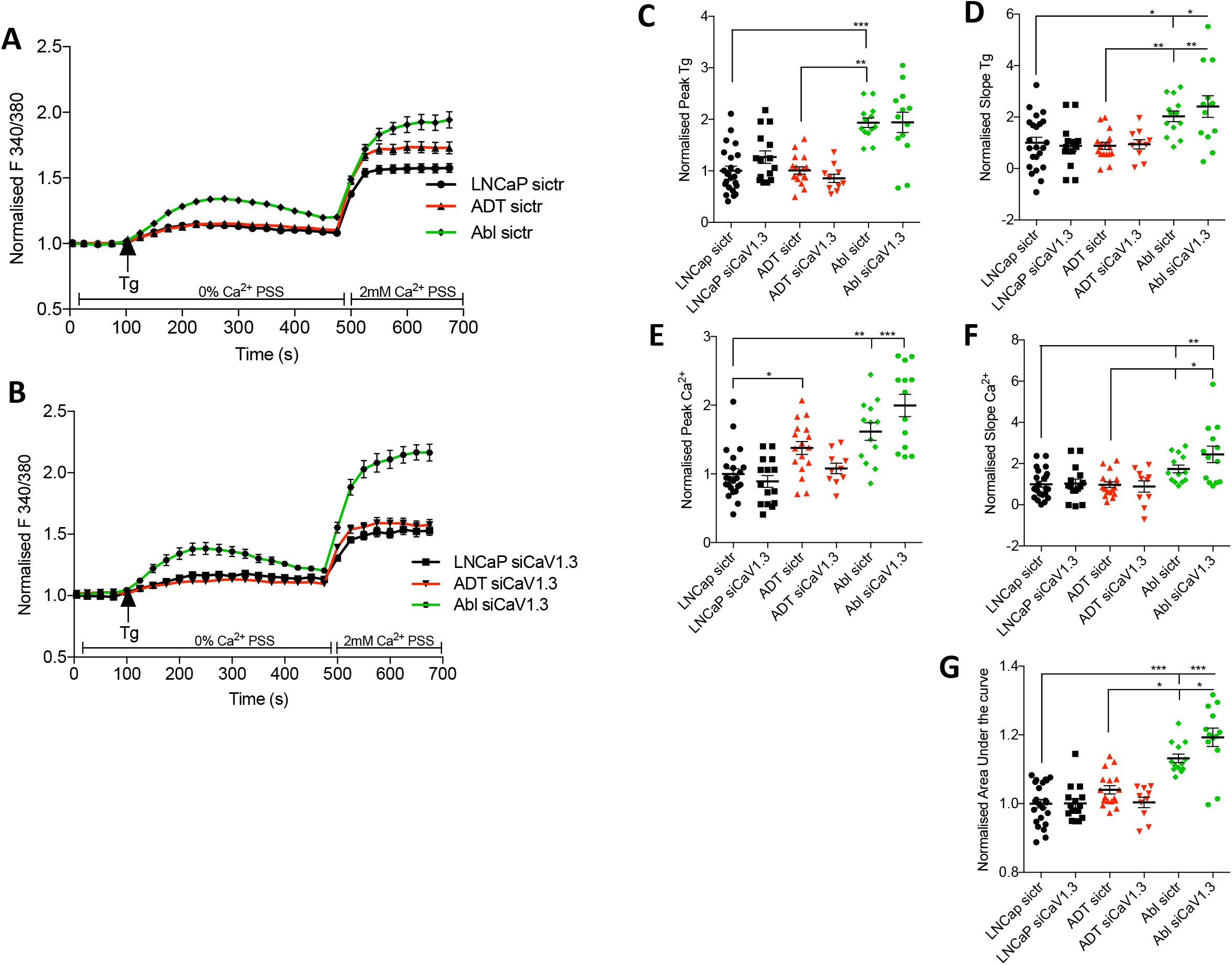
CaV1.3 drives aberrant SOCE following androgen deprivation. Fura-2am F340:380 ratiometric Ca^2+^ traces over time (s) displaying store release induced by Thapsigargin (Tg) in 0mM Ca^2+^ followed by store operated calcium entry in the presence of 2mM extracellular Ca^2+^ under treatment of **(A)** non-targeting control siRNA (sictr) in LNCaP (n=23), LNCAP-ADT(n=17) and LNCaP-Abl(n=13) or **(B)** siRNA targeting CaV1.3 (SiCaV1.3) in LNCaP (n=15), LNCAP-ADT(n=10) and LNCaP-Abl(n=13) (Two-way ANOVA, Dunnett’s MCT). Dot plots displaying the calculated **(C)** fold change Tg peak, (D) Tg slope, **(E)** fold change calcium peak, **(F)** calcium slope and **(G)** Area under the curve across each cell type and normalised to LNCaP sictr (Two-way ANOVA, Tukey’s MCT and t-test between sictr and sicav1.3).

Ca^2+^ entry following store release with Tg was measured after the addition of 2mM external Ca^2^. Here an increase in F340/380 ratio was observed under sictr conditions in LNCaP (1.58+/-0.03sem), LNCaP ADT (1.72+/-0.05) and LNCaP-Abl (1.94+/-0.06sem) (Figure 4A, all p<0.0001). Compared to LNCaP, ADT had a significant effect (P<0.0001) on peak Ca^2+^ entry producing a further 1.38 fold (+/-0.09sem, p<0.0408) increase in LNCaP ADT and a 1.62 fold (+/-0.13sem, p<0.0001) increase in LNCaP-Abl (Figure 4E). To confirm CaV1.3 contributed to this increase knockdown was employed. Here LNCaP, produced a similar F340/F380 under siCaV1.3 to sictr (Figure 4B). Under short term ADT CaV1.3 knockdown completely abolished the rise in Ca^2+^ entry observed in LNCaP-ADT, reducing it to 1.58+/-0.04sem (Figure 4B). Equating to a 29.83% decrease (p<0.0388) similar to the levels of Ca^2+^ entry witnessed in LNCaP (Figure 4F). Interestingly, CaV1.3 knockdown under long term ADT, increased Ca^2+^ entry by a further 37.97% (p<0.077) in LNCaP-Abl to 2.16+/-0.07sem F340/F380 (Figure 4B and E). Analysis of the Ca^2+^ entry slope found a similar effect in LNCaP-Abl cells only (Figure 4F), with an observed 1.74 fold increase (+/-0.19sem) under sictr and a further increase to 2.45 fold (+/-0.4sem) under siCaV1.3 (both p<0.0001).

Total change in SOC was determined through an area under the curve analysis (Figure 4G). This mirrored that observed with an increase in LNCaP-ADT (1.04+/-0.01sem) which was lost under siCaV1.3 (1.004+/0.02sem, p<0.0408), reducing down to similar levels observed with LNCaP. In LNCaP-Abl an increase of 1.13+/-0.01sem (p<0.0001) was observed, which was enhanced further under siCaV1.3 to 1.19+/-0.03sem (p<0.0001) (Figure 4G). In addition to this, we also noted that the calcium channel blocker (CCB), nifedipine, had no effect on SOCE unlike that observed with siCaV1.3 (Figure S4).

### CaV1.3 mediated increase in SOCE promotes cell proliferation and survival

Final investigations where conducted to determine the impact of CaV1.3 mediated SOCE on PCa biology under ADT. PCa proliferation was found to be reduced under short term ADT in LNCaP-ADT (0.56FC+/-0.07sem, p<0.0003). Whereas proliferation of the long term ADT androgen insensitive LNCaP-Abl cells increased 1.73FC+/-0.1sem (p<0.0001) (Figure 5A). Knockdown of CaV1.3 had no effect on the proliferation of LNCaP or LNCaP-ADT cells but did produce a 39.59% reduction in LNCaP-Abl (+/-3.1sem, p<0.0001) (Figure 5Bi-iii). Cell survival measured by colony forming assay (Figure 5C) found LNCaP averaged 48 colonies (+/-8.8sem), which was reduced under short term ADT to 37 colonies (+/-2.9sem). However, LNCaP-Abl displayed an increase in cell survival producing 167 colonies (+/-6.9sem, p<0.05), resulting in a 3.5 fold increase. Under SiCaV1.3 no effect was witnessed on cell survival of LNCaP or LNCaP-ADT (Figure Di + ii), however LNCaP-ABL displayed a colony reduction of 82.8% (+/-6.3sem, p<0.029,Figure 5Diii). Lastly, metastatic potential was measured (Figure 5Ei) with LNCaP cells demonstrating the highest cell index score (0.09+/-0.07sem), which was reduced by 155% in LNCaP-ADT and 223% in LNCaP-Abl (p<0.0028)(Figure 5Eii). Under the same conditions CaV1.3 knockdown was observed to have no detectable effect.

**Figure 5:**
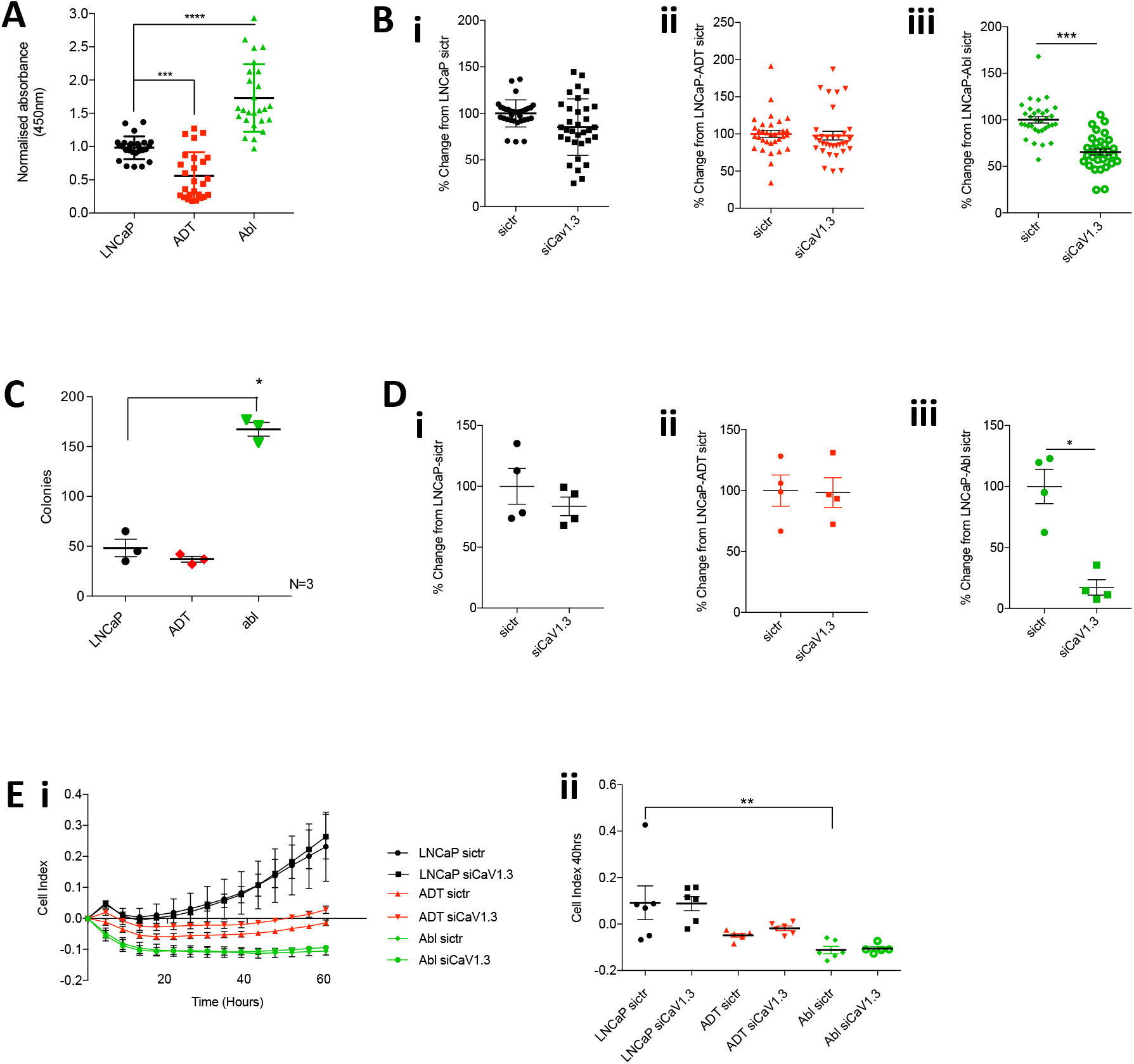
CaV1.3 mediated aberrant SOCE promotes cell proliferation and survival under long term ADT. Cell proliferation of LNCaP, LNCaP ADT and LNCaP Abl **(A)** was measured by WST-1 assay with fold change calculated and represented in a dot plot normalised to LNCaP (One-way ANOVA, Tukey MCT, N=9, n=26). **(B)** Fold change in cell proliferation was determined for **(i)** LNCaP, **(ii)** LNCaP-ADT, and **(iii)** LNCaP-Abl, treated with either siRNA control (sictr) or CaV1.3 siRNA (siCaV1.3) (unpaired t-test, N=12, n=33). Cell survival capacity was measured by colony formation assay **(C)** Total colonies counted between LNCaP, LNCaP ADT or LNCaP abl (One-way ANOVA, Tukey MCT, N=3). **(D)** Fold change in colony count after treatment with sictr or siCaV1.3 for **(i)** LNCaP, **(ii)** LNCaP-ADT, and **(iii)** LNCaP-ABL (Mann-Whitney t-test, N=4). (**Ei)** Migration and invasion capacity was measued in CIM plates as cell index over the course of 60 hours in PC cells treated either with sictr or siCaV1.3 (N=6), with **(Eii)** measurements taken at 40hours for analysis between cells and treatments (Kruskal-Wallis MCT, N=6).

## Discussion

Multiple studies have demonstrated that the expression of the L-type VGCC, *CACNA1D*/CaV1.3, is positively correlated with progression to CRPC, yet the functional role of this channel and its impact on cancer biology under ADT has remained unknown until now. We sought to investigate these aspects and in particular focus on how the channel contributes to ADT resistance leading to CRPC. Here we demonstrate for the first time that CaV1.3 represents an oncogenic mechanism that drives ADT resistance by promoting aberrant SOCE through a novel non-canonical mechanism, which enhances cell proliferation and survival in CRPC.

Initial bioinformatic evaluation of *CACNA1D* expression in PCa patient samples highlighted an upregulation in primary PCa which was further increased in metastatic CRPC, matching similar previous studies^10,13,15^. In addition, this work also supports other studies linking channel expression to increased Gleason score^13,30^. Owing to the fact that ADT resistance promotes CRPC^31^, we investigated if channel expression increased during ADT. This was confirmed through the observed increase of *CACNA1D* expression in ADT-treated vs non-treated group. This has never been shown previously and suggests upregulated *CACNA1D* drives ADT resistance. Furthermore, as a known ERG target previous studies have also demonstrated a correlation between *CACNA1D* expression and positive TMPRSS2:ERG fusion gene status^15,26–29,32^. Interestingly, we witnessed this effect in primary PCa only. However, in metastatic CRPC and ADT treated samples, *CACNA1D* expression was elevated regardless of ERG status. This novel observation demonstrates that increased CANCA1D expression can also occur in TMPRSS2:ERG negative patients treated with ADT.

To further validate the link between ADT and CaV1.3 expression, a LNCaP xenograft mouse and cell line model were employed with both displaying a significant increase in CaV1.3 expression under ADT compared to untreated control. Studies using this mouse model have previously demonstrated that bicalutamide caused vascular collapse and increased hypoxia in these tumours, exerting a selective pressure which favours growth of treatment-resistant cells^23^. Similarly, the mouse tumour and PCa cell line analysis presented here now suggests a link between increased CaV1.3 and ADT resistance, which also corroborates our observations in patient samples. While this is the first study to demonstrate that ADT promotes CaV1.3 expression in PCa, other research confirms that this treatment can influence expression of calcium channel families including T-Type VGCC, CaV3.2^33,34^.

This initial work highlighted the suitability of the outlined cell model for further investigations into the channels ability to modulate Ca_i_^2+^ under ADT and associated impact on cancer biology, which to our knowledge had not been studied. As a VGCC, CaV1.3 is known to mediate Ca^2+^ influx from the plasma membrane upon membrane depolarisation^35^. However, the addition of high external potassium solution failed to induce any changes in Ca_i_^2+^ either with and without ADT. This lack of traditional canonical function can be explained by the fact that the full length 250kDa isoform, which is known to work through this mechanism^18^, was found expressed at low levels in the cytosol. This idea was also corroborated by our preliminary patch clamp data were no detectable L-type VGCC calcium influx was found on the plasma membrane with or without androgens present (Data not shown). Further investigations into Ca_i_^2+^ under ADT in our cell model found an increase in basal Ca_c_^2+^ under short and long term treatment, which suggested a potential link to store operated calcium.

To explore this further we first confirmed the presence of SOCE in our PCa cell line model and found that it increased with ADT and duration of treatment. A number of other studies have also demonstrated the presence of SOCE in a range of PCa cell lines^36–39^ including LNCaPs used here in. However, in contrast to our work a decrease in SOCE was observed following ADT ^21,36,40^. This variation appears to be related to fact that these studies use shorter treatment time frames and induce androgen deprivation artificially with charcoal stripped media, known to have wide spread cellular effects^41^. In contrast our cellular model used a more clinically relevant treatment with bicalutamide, which specifically blocks androgen signalling only. In addition, this research and others highlight that longer term ADT leads to further changes at the cellular and gene level not found with short term treatment^42–44^.

To assess the impact of increased CaV1.3 on the observed aberrant SOC under ADT an siRNA approach was employed. Interestingly, CaV1.3 knockdown completely abolished the increase in SOCE under short term ADT, while the long-term ADT CRPC LNCaP-Abl cell line observed a further increase in SOCE. This is the first time that CaV1.3 has been shown to influence SOCE in PCa but the underlying mechanism is not clear. Research in other tissues does show coupling of CaV1.3 to intracellular stores through interaction with NCX^19^ and also RYR^18,45–47^. In particular it has been suggested that this interaction is isoform specific^18^, which would correlate with the presence and upregulation of the shorter 170kDA CaV1.3 in our cell model. Furthermore, this shorter isoform is known to lack sensitivity to CCB’s^18^, explaining why the CCB nifedipine failed to impact on SOCE. This important observation potentially accounts for the conflicting epidemiology reports investigating the beneficial effects of CCB use in PCa^48–51^. Overall this suggests that CaV1.3 is mediating SOCE via a non-canonical mechanism through the interaction with other store release channels. Alternatively, CaV1.3 has been previously shown to regulate gene transcription through its cleaved 70kDa c-terminus product^52,53^. This study showed the presence of the CaV1.3 c-terminus in the nucleus under short term ADT which was lost under long term treatment and progression to CRPC (Figure S1A). This difference could account for the opposing effects on CaV1.3 knockdown and potentially suggests that the c-terminus could be playing a role influencing the expression of calcium channels involved in the modulation of SOCE.

As outlined aberrant SOCE is known to contribute to various cancer hallmarks, promoting treatment resistant and disease progression^8^. Thus the study sought to investigate the impact of CaV1.3 mediated SOCE on PCa biology under ADT. As expected and observed by other studies^42,44^, short term ADT reduced proliferation, colony formation and migration. Knockdown of CaV1.3 under these conditions had no effect, suggesting that the channel does not affect these aspects of PCa biology at this stage but further investigations into other cancer hallmarks are required. In contrast, under long term ADT, cell proliferation and colony formation was increased. This is indicative of progression to an androgen-independent castrate-resistance state as witnessed in patients^54^ and shown in PCa cell lines such as LNCaP-Abl^22,55,56^. A number of mechanisms have been proposed to promote this phenotype such as AR alternations^57^, however changes in Ca_i_^2+^ such as SOCE have also been linked to PCa progression^37,58,59^. This suggests that the observed increase in SOCE through CaV1.3 in LNCaP-Abl is promoting ADT resistance and contributing to the increase in proliferation and colony formation. Furthermore, the decrease in proliferation and CFA after CaV1.3 knockdown in our long term ADT model could be explained by the dual role of Ca_i_^2+^ modulation where a sustained or large Ca_i_^2+^ increase such as that observed with our SOCE is known to induce apoptosis^37,60^.

### Conclusion

Overall this novel study demonstrates for the first time that upregulated CaV1.3 contributes to ADT resistance by driving aberrant SOCE through a non-canonical mechanism which promotes CRPC biology. Bioinformatic analysis highlighted an associated increase in *CACNA1D* expression with ADT and Gleason score. In addition, while the *TMPRSS2:ERG* mutation is a known driver of *CACNA1D* expression, ADT can also result in similar increases in expression in *TMPRSS2:ERG* negative patients. This highlights the channel as a potential biomarker for PCa disease progression to CRPC in all patients who are receiving ADT. This study also provides new evidence that CaV1.3 function is determined by the isoforms expressed, as we demonstrate that in PCa the 170kDA CaV1.3 mediates SOCE. Moreover, the subsequent increase in CaV1.3 SOCE plays a key role in the development of resistance to ADT enabling PCa biology that contributes to CRPC. Of note, CCB’s do not influence Ca_i_^2+^ through this isoform, explaining the conflicting studies investigating the impact of CCB use in PCa. However, this research does highlight that targeting different CaV1.3 isoforms and or other SOC channels could be of therapeutic benefit to later stage ADT resistant disease where there is an unmet need for new treatments. This exciting potential for patient benefit means that further research on CaV1.3 isoforms and associated interacting partners mediating SOCE in PCa is highly recommended.

## Abbreviations

(ADT): Androgen Deprivation Therapy
(AR): Androgen Receptor
(Ca^2+^): Calcium
(CCB): Calcium Channel Blocker
(CRPC): Castrate Resistant Prostate Cancer
(CIM): Cellular Invasion and Migration
(CFA): Colony Formation Assay
(Ca_c_^2+^): Cytosolic Calcium
(GAPDH): Glyceraldehyde 3-phosphate dehydrogenase
(IF): Immunofluorescence
(Ca_i_^2+^): Intracellular Calcium
(MCT): Multiple Comparison Test
(PCa): Prostate Cancer
(Tg): Thapsigargin
(TMA): Tumour Microarrays
(SiCaV1.3): siRNA CaV1.3
(sictr): SiRNA Control
(SOC): Store Operated Current
(SOCE): Store Operated Calcium Entry
(VGCC): Voltage Gated Calcium Channels

