## Supplemental Figures and Tables for "CaV1.3 enhanced store operated calcium promotes resistance to androgen deprivation in prostate cancer"

### Supplementary Table 1

A

| Origin | ADJACENT | MET | PRIMARY |
|---|---|---|---|
| Asian | 1 | 0 | 2 |
| Black Hispanic | 0 | 0 | 3 |
| Black Non-Hispanic | 0 | 1 | 24 |
| Unknown | 2 | 0 | 4 |
| White Hispanic | 0 | 2 | 0 |
| White Non-Hispanic | 25 | 15 | 97 |

B

| Hormone therapy | Tumour Type | | | Total |
|---|---|---|---|---|
| | Adjacent | Metastatic | Primary | |
| Post-ADT | 0 | 7 | 3 | 10 |
| ADT | 2 | 8 | 16 | 24 |
| No Treatment | 26 | 3 | 111 | 114 |

C

| Gleason Score | ADJACENT | MET | PRIMARY |
|---|---|---|---|
| 6 | 12 | 0 | 41 |
| 7 | 13 | 2 | 73 |
| 8 | 1 | 3 | 8 |
| 9 | 1 | 4 | 7 |
| No data | 1 | 9 | 1 |

D

| ERG Status | ADJACENT | MET | PRIMARY |
|---|---|---|---|
| Negative | 19 | 14 | 78 |
| Positive | 3 | 4 | 30 |
| Flat | 6 | 0 | 22 |

E

| Hormone therapy | ERG status | Totals |
|---|---|---|
| Post-ADT/ADT | Negative | 23 |
| | Positive | 9 |
| | Flat | 2 |
| | Subtotal | 34 |
| No treatment | Negative | 70 |
| | Positive | 26 |
| | Flat | 18 |
| | Subtotal | 114 |
| Total | | 148 |

***Supplementary Table 1: Sample details used for bioinformatic analysis (A)*** *Tumour type vs origin. This sample set contained expression information from a total of N=176 samples of which n=130 where primary cancers and n=18 metastatic. A subset of primary cancers also had n=28 paired adjacent normal samples.* ***(B)*** *Sample size and tumour type for each hormone therapy group used.* ***(C)*** *Sample source and Combined Gleason score showed that most samples had scores of 6 or 7 than 8 or 9.* ***(D)*** *Sample source and ERG status showed that most were ERG-negative (73%) and a minority were ERG-positive (27%) and that these proportions were similar in both metastatic (78% and 22%) and primary tumours (72% and 28%).* ***(E)*** *Cohort ADT hormone therapy treatment and ERG status numbers.*

### Supplementary Figure 1 – CaV1.3 location

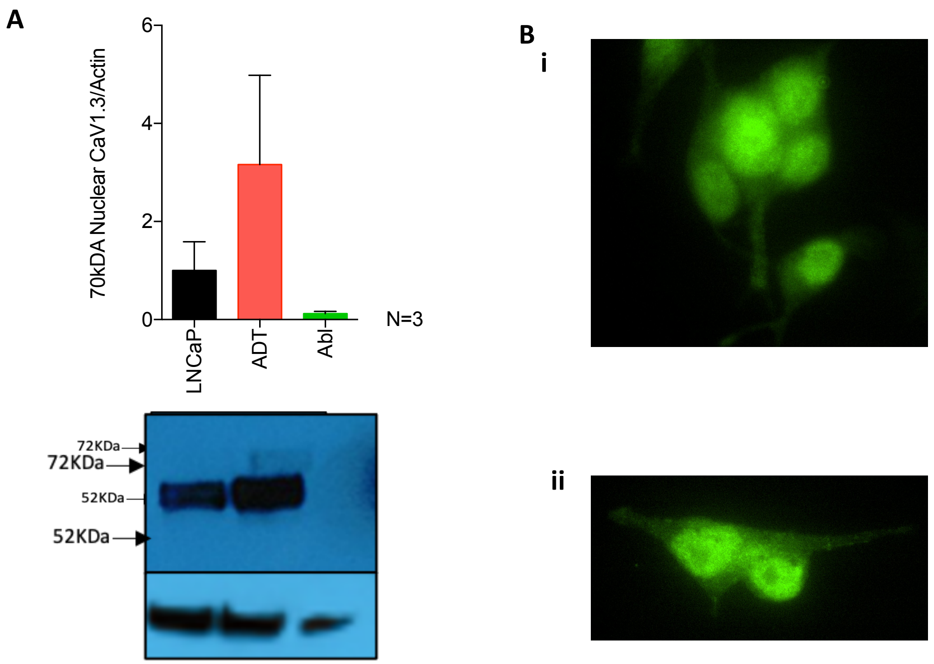


***Figure S1: CaV1.3 cellular location.*** **(A)** protein fractionation blot showing a 60kDa isoform expressed in the nucleus of both LNCaP and LNCaP-ADT cells. **(B)** immunofluorescence images of CAV1.3 stained **(i)** LNCaP and **(ii)** LNCaP-ADT cells demonstrated a mainly intracellular and nuclear location.

.

### Supplementary Figure 2 – Osmolarity controls for high potassium tests

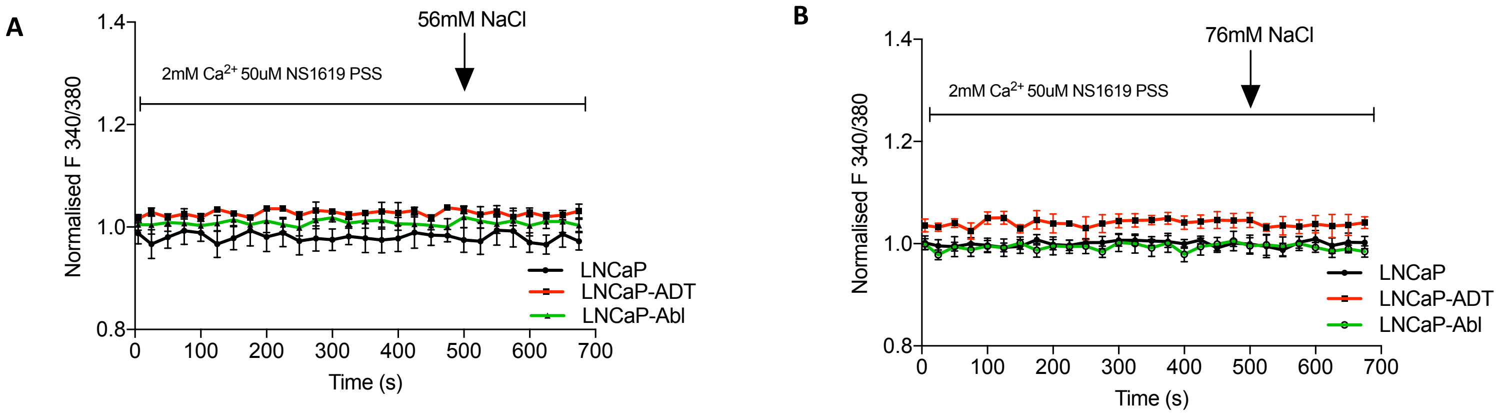


***Figure S2: NaCl osmolarity control for high potassium test.*** *Fura-2am calcium traces over time of osmolarity controls where androgen sensitive LNCaP cells, LNCaP-ADT cells treated with 10µM bicalutamide (7-10 days) and androgen insensitive long-term androgen deprived LNCaP-Abl cells where depolarised with high external sodium concentrations of* ***(A)*** *56mM NaCl or* ***(B)*** *76mM NaCl.*

### Supplementary Figure 3– Validation of CaV1.3 KD

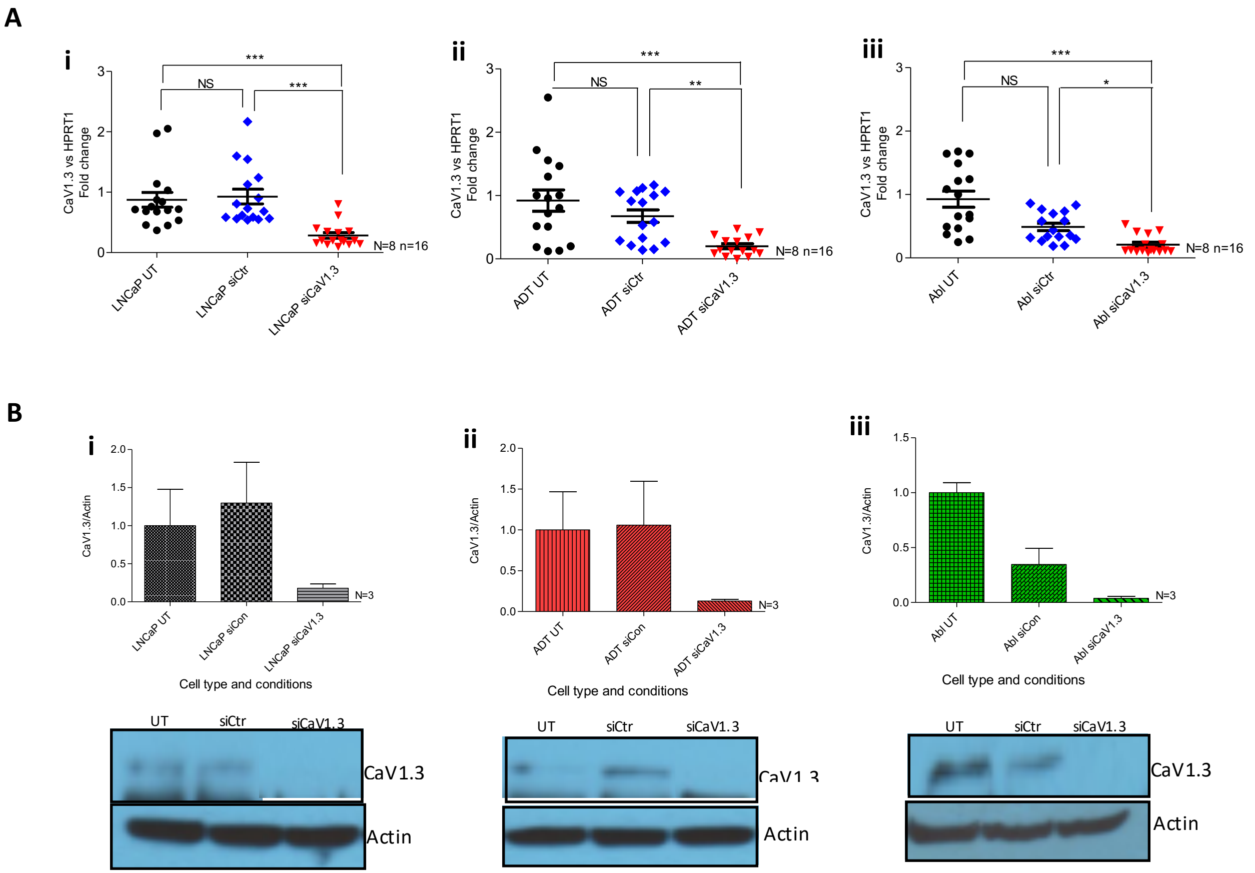


***Figure S3: Gene and protein expression of CaV1.3 after knockdown. (A)*** *PCR analysis of expression of CaV1.3-fold change from HPRT1 in (i) LNCaP, (ii) LNCaP-ADT, (iii) LNCaP-Abl cells transfected with control siRNA (blue) or siRNA targeting CaV1.3 (red), normalised to untransfected cell of type (black) (Kruskil-Wallis, Dunn's MCT, N=8, n=16).* ***(B)*** *Western blot analysis of expression of CaV1.3 fold change from Actin in (i) LNCaP, (ii) LNCaP-ADT, (iii) LNCaP-Abl cells transfected with control siRNA or siRNA targeting CaV1.3, normalised to untransfected cell of type (black) (Kruskil-Wallis, Dunn's MCT, N=3). * P<0.05, ** P<0.01, ***P<0.001, NS not significant.*

### Supplementary Figure 4- Calcium channel blockers have no effect on SOC calcium

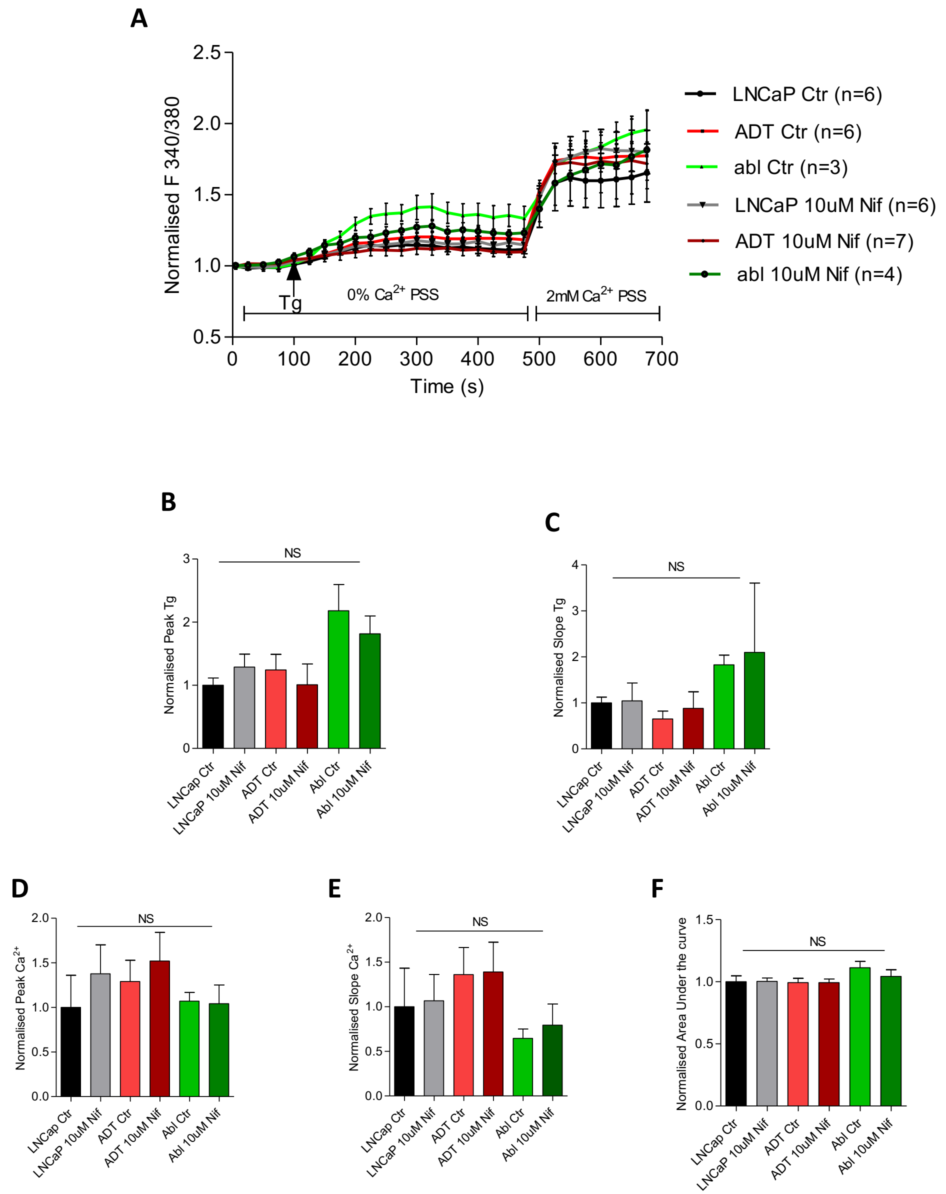


***Figure S4: Nifedipine has no significant effect on the $Ca^{2+}$ mobilisation in LNCaP cells at any stage under ADT**: **(A)** Store operated calcium determined by Fura 2-AM ratio metric analysis of calcium concentration over time(s). Analysis of androgen sensitive LNCaP cells , LNCaP ADT LNCaP Abl treated with either 10µM Nifedipine or DMSO control. Dot plots displaying the calculated normalised **(B)** Tg peak, **(C)** Tg slope, **(D)** calcium peak, **(E)** calcium slope and **(F)** Area under the curve. Analysed using Kruskil-Wallis significance test between cell types and treatments.*
